# Diagnosis of Duchenne Muscular Dystrophy using Raman Hyperspectroscopy

**DOI:** 10.1101/2020.01.08.897793

**Authors:** Nicole M. Ralbovsky, Paromita Dey, Andrew Galfano, Bijan K. Dey, Igor K. Lednev

**Author notes:** Corresponding authors: Bijan K. Dey, Ph.D., and Igor K. Lednev, Ph.D.,.

## Abstract

Duchenne muscular dystrophy (DMD) is the most common and severe form of muscular dystrophy and affects boys in infancy or early childhood. DMD is known to trigger progressive muscle weakness due to skeletal muscle degeneration and ultimately causes death. There are limited treatment regimens available that can either slow or stop the progression of DMD. An accurate and specific method for diagnosing DMD in its earliest stages is needed to prevent progressive muscle degeneration and death. Current methods for diagnosing DMD are often laborious, expensive, invasive, and typically diagnose the disease later on it is progression. In an effort to improve the accuracy and ease of diagnosis, this study focused on developing a novel method for diagnosing DMD which combines Raman hyperspectroscopic analysis of blood serum with advanced statistical analysis. Partial Least Squares Discriminant Analysis (PLS-DA), was applied to the spectral dataset acquired from control and *mdx* blood serum of 3- and 12-month old mice to build a diagnostic algorithm. Internal cross-validation showed 95.2% sensitivity and 94.6% specificity for identifying diseased spectra. These results were verified using external validation, which achieved 100% successful classification efficiency at the level of individual donor. This proof-of-concept study presents Raman hyperspectroscopic analysis of blood serum as a fast, non-expensive, minimally invasive and early detection method for the diagnosis of Duchenne muscular dystrophy.

## Introduction

Duchenne muscular dystrophy (DMD) is a progressive form of muscular dystrophy which typically affects male infants. DMD is an X-chromosome linked recessive disorder caused by a mutation of the dystrophin gene, which results in progressive weakness and atrophy of the skeletal and heart muscles.^1,2^ Symptoms can begin in boys as young as 1 to 6 years old, and initially include difficulty sitting, standing, walking or speaking.^3^ The issues associated with DMD are severe, worsen overtime, and greatly impact the well-being of the afflicted individual. In fact, secondary complications due to DMD, including heart and respiratory muscle problems, can lead to life-threatening conditions.^4^ Limited treatments exist for DMD, which can stop the progression of the disease and help control the symptoms associated with it.

Diagnosing DMD typically involves evaluating family history as well as conducting blood tests to assess the levels of specific muscle enzymes in the blood. Although the inheritance of the disease is through an X-linked recessive pattern, there are cases where DMD occurs in families who have no history of it. The complicated pattern of inheriting DMD suggests a need for additional testing. Blood tests often monitor the level of serum creatine phosphokinase (CPK), with high levels indicating muscle damage is causing the muscle weakness. However, this test can only detect the disease in later stages and is generally non-specific, as high levels of CPK can be found in an individual’s blood after experiencing a heart attack, drinking alcohol in excess, or participating in strenuous exercise.^5–10^ Electromyography is often used to confirm muscle weakness without pinpointing a direct cause of it.^11^ Muscle biopsies can differentiate muscular dystrophies from other muscle diseases^12^, however biopsy examinations can be both expensive and invasive for the individual undergoing testing. Genetic testing can confirm if there is a mutation within the DMD-causing gene, as well as distinguish between different types of muscular dystrophy. However, because genetic testing and muscle biopsies are invasive and expensive, these options are typically pursued only after other options have been exhausted, thus resulting in the disease being diagnosed in its later stages. Because DMD is progressive, and its symptoms worsen overtime if treatment isn’t initiated, it is of the utmost importance to definitively diagnose the disease as early on in its progression as possible, before symptoms become too severe. The earlier the disease is identified within its progression, the better opportunity the afflicted individual has for seeking effective treatment opportunities.

To improve the accuracy and ease and potential of an early diagnosis, we focused on developing a novel method for diagnosing DMD using Raman hyperspectroscopic analysis of *mdx* mouse blood serum combined with advanced statistical analysis. The dystrophin mutant *mdx* mice do not express dystrophin and have been widely used as a model system to study DMD and to make important advances in understanding therapeutic strategies; it has allowed for the molecular processes and underlying causes of the disease to be better understood.^2,13^ The *mdx* mouse model serves as an efficient and useful model for developing a better diagnostic method without influence from complications, such as the effect of prescribed medications, associated with humans.

Raman hyperspectroscopy has shown it has great potential to diagnose many diseases including cancers,^14,15^ Alzheimer’s disease,^16–18^ and other diseases where pathophysiological changes occur.^19,20^ Raman hyperspectroscopy involves collecting multiple Raman spectra from a sample to better characterize its inherent heterogeneity. This generates a three dimensional data cube (*x, y, λ*) where the *x* and *y* dimensions correspond to spatial coordinates and the *λ* dimension represents the Raman spectrum collected at a particular pair of coordinates. Through collection of multiple spectra per sample, its biochemical composition is better understood; as such, a change in biological composition of blood serum due to disease progression can be detected using Raman hyperspectroscopy. This technique produces a specific spectral fingerprint which represents the biochemical composition of the sample analyzed. This specific information can thus be used to distinguish between different samples, such as body fluids collected from healthy donors and from donors with a disease. Here, we capitalized on the advantages of Raman hyperspectroscopy in combination with advanced statistical analysis to build a model which identifies spectral differences between different classes of samples to make diagnostic predictions. Partial Least Squares Discriminant Analysis (PLS-DA) was used to build a model which could distinguish Raman spectral data of healthy control mice from Raman spectral data of *mdx* mice. The results were verified using external cross-validation. Genetic Algorithm (GA) was then used to identify the spectral features which contribute the most useful information toward differentiation. The spectral features identified by GA were assigned to vibrational modes of various biomolecules which were previously identified as playing a role in the pathogenesis of DMD. For the first time, this proof-of-concept study shows Raman hyperspectroscopy in combination with advanced statistical analysis is successful in detecting DMD in a simple, accurate, early, and minimally invasive manner.

## Results

### Validation of skeletal muscle abnormalities in *mdx* mice by examining the Tibialis Anterior (TA) muscle morphology

Duchenne muscular dystrophy is the most common and most severe form of muscular dystrophy. DMD is characterized by muscle wasting and weakness due to excessive muscle degeneration. The Tibialis Anterior (TA) muscle morphology of 3-month old and 12-month old control (C57BL/10ScSnJ) and *mdx* (C57BL/10ScSn-Dmd<mdx>/J) mice was examined using Hematoxylin and Eosin (H&E) staining (Figure 1 A-D). As expected, normal skeletal muscle morphology was observed in 3-month old control mice (Figure 1A). Mild skeletal muscle degeneration was observed in 3-month old *mdx* mice as characterized by the smaller diameter of muscle fibers with central nuclei, occasional presence of atrophied muscle fiber, and the presence of an increased number of nuclei representing inflammatory cells (Figure 1B). Similar to 3-month old control mice, 12-month old control mice displayed normal skeletal muscle morphology (Figure 1C). Skeletal muscle degeneration progresses as *mdx* mice get older. As such, muscle degeneration was much more prominent in the 12-month old *mdx* mice as marked by the absence of normal muscle structure in most areas of the tissue section and the presence of fatty and fibrotic tissues (Figure 1D).

**Figure 1.**
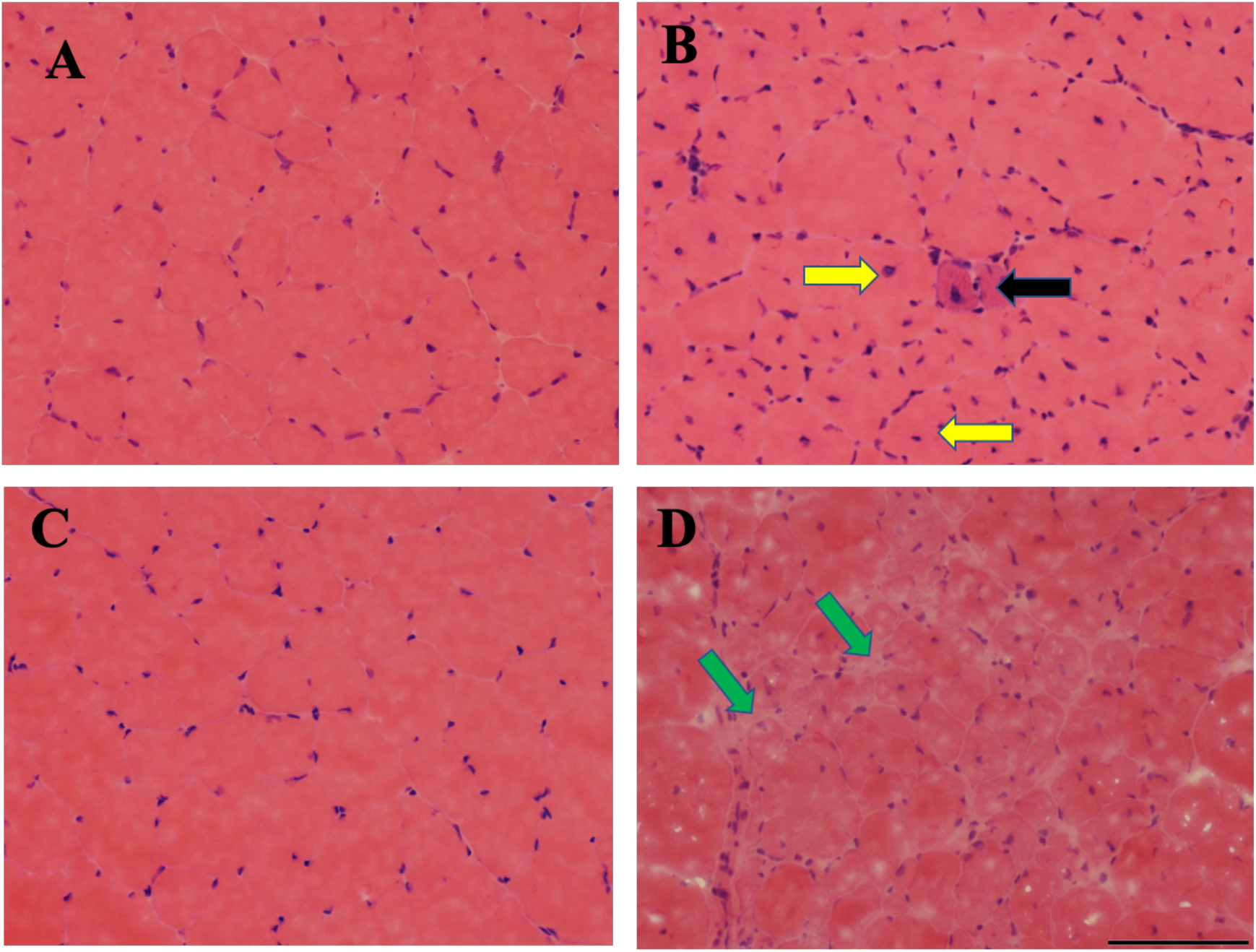
Skeletal muscle degeneration is observed in the mouse model of DMD. Hematoxylin and Eosin (H&E) staining of TA muscle cross sections from 3- and 12-month-old control (C57BL/10ScSnJ) (A, C) and *mdx* (C57BL/10ScSn-Dmd<mdx>/J) (B, D) mice. The 3-month old control muscle cross-section shows normal morphology (A) whereas 3-month old *mdx* mice show muscle degeneration (denoted by muscle with central nuclei and smaller diameter, yellow arrows, atrophied muscle, black arrow, and more prevalent nuclei representing inflammatory cells) (B). Control mice at 12-months old (C) are compared to the 12-month old *mdx* mice (D) where muscle degeneration is much more dramatic, as evident by the absence of normal muscle structure in almost all areas of the section; the muscle structure is often taken over by fatty and fibrotic tissues, as indicated by green arrows. Scale Bar: 100 uM.

### Raman spectroscopic analysis of mice blood serum

Because DMD is both progressive and treatable, it is crucial to diagnose the disease as early as possible. In this proof-of-concept study, blood serum of healthy and *mdx* mice at 3- and 12-months old was analyzed by Raman hyperspectroscopy in an attempt to develop a novel diagnostic method. Blood serum is the portion of blood which does not contain cells or clotting factors, and has been widely studied in the past for diagnostic purposes.^17,21–24^ Only 10 μL of blood serum was required from each donor. The serum was deposited on an aluminum substrate and allowed to dry before conducting Raman hyperspectroscopic analysis.

Raman spectra were collected from the serum of 14 mice donors through automatic mapping. Mapping was conducted to obtain an accurate representation of the entire biochemical composition of each sample, with the intention of identifying key biochemical components useful for discrimination between classes. The two classes of donors consisted of healthy mice (control, n=7) and *mdx* mice (MDX, n=7). Of the 14 total blood serum samples, six (three control and three MDX) were collected from mice at three months old and eight (four control and four MDX) were collected from mice at 12 months old. Different ages of mice were used in order to illustrate the method’s ability to detect the disease early on in its progression. The mean preprocessed spectra for all donors from each class is seen in Figure 2.

**Figure 2.**
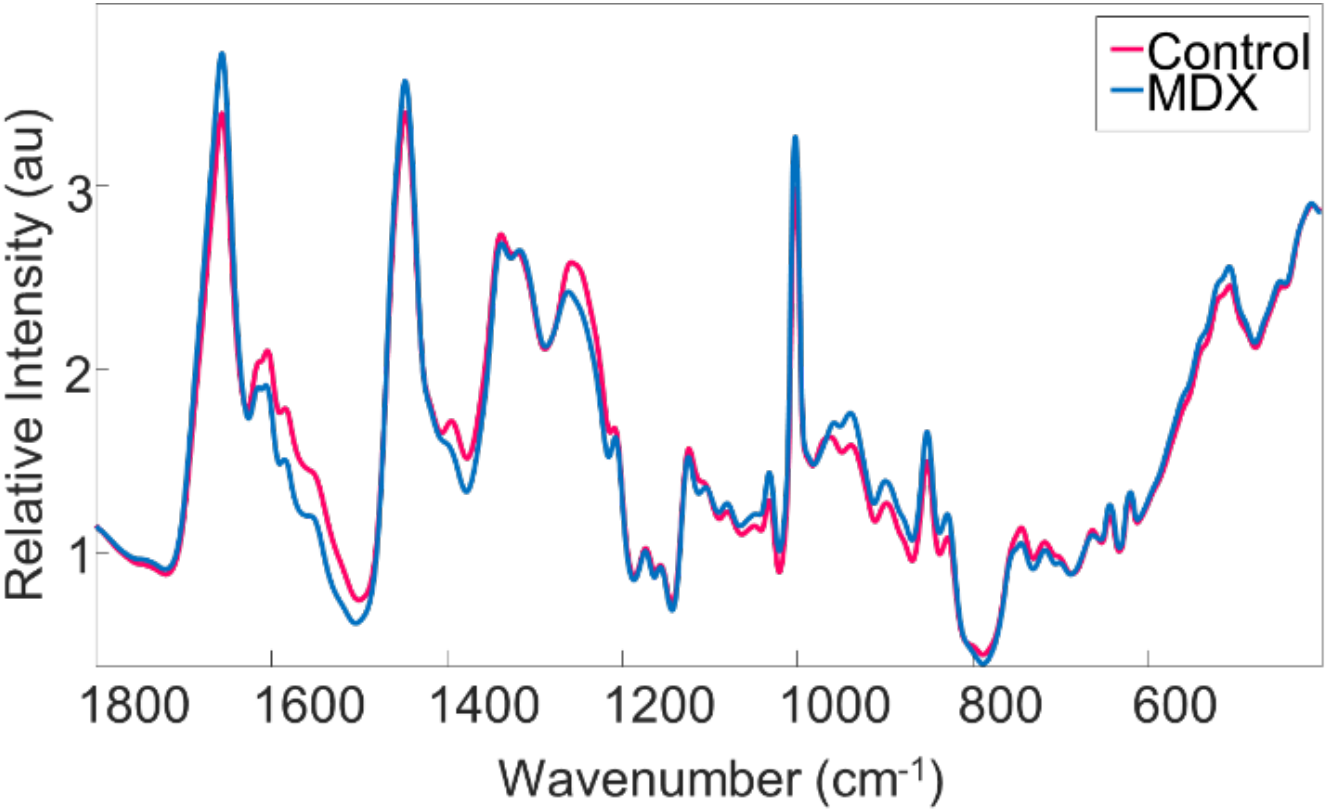
Mean Raman spectra collected from the two classes of mice blood serum. The mean spectrum of all control mice blood serum samples is represented by the pink line, whereas the mean spectrum of all *mdx* mice blood serum samples is represented by the blue line.

### Model calibration for differentiating healthy controls from MDX mice

The donors were split into two groups: the calibration group and the validation group. Ten of the donors were used in the calibration set (five control, five MDX); the spectral data from these donors was used to build the PLS-DA prediction algorithm. The validation dataset, consisting of spectral data from two control donors and two MDX donors, was used for external validation. Mice of different ages (3- and 12-months) were included in both the calibration and validation groups.

The difference between the mean control spectrum and the mean MDX spectrum was calculated and compared with ±2 standard deviations within each class. It was observed that the difference spectrum fell within the standard deviations (Supplementary information, Figure S.1). This indicates that the spectral changes shown in the difference spectrum (Figure S.1) are smaller than the variation which occurs within each class, and thus are statistically insignificant. As such, advanced statistical analysis was required to capitalize on the important spectral features which vary between the two classes at the level of individual spectra but are hidden from the mean spectra. This variability is useful for discriminating between the two classes of data.

To uncover the differences between individual spectra to be used for diagnostic purposes, Partial Least Squares Discriminant Analysis (PLS-DA) was selected to build a discrimination algorithm. A binary model was built to distinguish between control and MDX blood serum spectral data of the calibration dataset. Eight latent variables were used to capture the maximum covariance between the spectral data and the assigned classes. Each spectrum from the calibration dataset was assigned a set of scores which correspond to how similar that spectrum is to each latent variable. Each class is thus ideally represented by a range of scores seen as typical for that class. Scores plots can be used to understand the separation which exists between different classes, and any spectrum which is loaded into the model will be given a set of scores which is used to decide to which class it belongs. The model built herein showed clear separation between the two classes (Figure 3).

**Figure 3.**
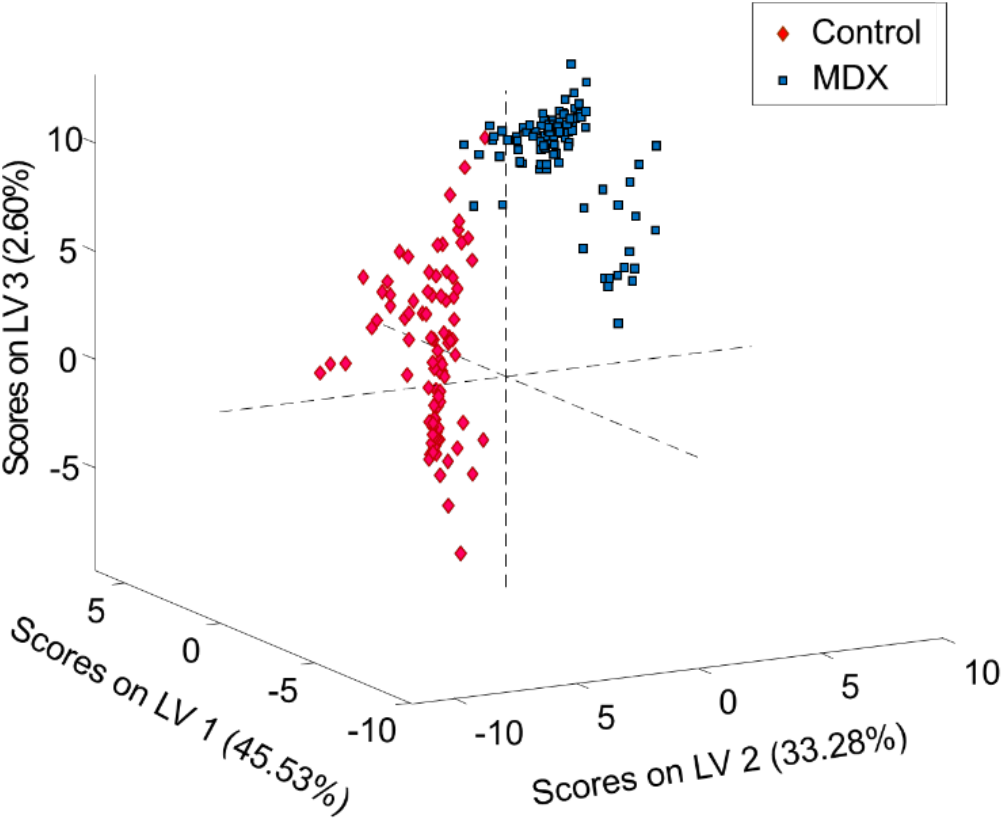
PLS-DA scores plot. The PLS-DA scores plot built using the first three latent variables. The distribution of symbols represents the separation which exists between the two classes of blood serum spectra where pink diamonds signify controls and blue squares signify MDX. Each symbol represents an individual spectrum.

The sensitivity and specificity rate for classification of the PLS-DA diagnostic algorithm were calculated. In this study, the sensitivity is defined as the true positive rate, or the percentage of MDX spectra correctly predicted as belonging to the MDX class. The specificity is defined as the true negative rate, or the percentage of control spectra correctly predicted as not belonging to the MDX class. Individual spectral predictions for all donors within the calibration dataset can be observed in the confusion matrix presented in Table 1. Here, every Raman spectrum is assigned a class (either control or MDX). The assignments are compared to the true, or known, classification for each spectrum. Internal cross-validation of the PLS-DA model by venetian blinds resulted in 95.2% sensitivity and 94.6% specificity for training the algorithm using the calibration dataset.

**Table 1.** Confusion matrix illustrating individual spectral predictions for all spectra of the calibration dataset

| Confusion Matrix – Internal Cross Validation |  |  |
| --- | --- | --- |
|  | Actual Class |  |
|  | Control | MDX |
| Predicted as Control | 212 | 11 |
| Predicted as MDX | 12 | 217 |

### External validation of the PLS-DA model

External validation was performed using the spectral data collected from the four donors of the validation dataset. The validation dataset was kept independent from the training set and is thus considered a powerful method for testing the validity and strength of the classification model as well as its potential for real-world applications. A total of 185 spectra collected from the four samples were loaded into the PLS-DA algorithm for external validation. The class assignment for each spectrum was predicted (Table 2). Once again, the sensitivity (true-positive rate) and specificity (true-negative rate) of classification for external validation were calculated. Here, 100% sensitivity and 87.0% specificity was achieved for external validation, calculated using the validation dataset.

**Table 2.** Confusion matrix illustrating individual spectral predictions for all spectra of the validation dataset

| Confusion Matrix – Internal Cross Validation |  |  |
| --- | --- | --- |
|  | Actual Class |  |
|  | Control | MDX |
| Predicted as Control | 80 | 0 |
| Predicted as MDX | 12 | 93 |

### Receiver Operating Curve analysis of external validation results

To better understand individual spectral predictions, a receiver operating characteristic (ROC) curve was used to identify the most optimum threshold for determining donor-level classifications based on spectral-level predictions. A ROC curve is typically used to evaluate the performance of a binary classifier, such as the PLS-DA model built here. The curve is generated by plotting true positive rate values (sensitivity) against false positive rates values (1-specificty); every point in the ROC curve corresponds to a potential threshold for discrimination. The most ideal threshold would exist at (0,1), which represents no false positives and a 100% true positive rate for predictions made at the donor-level. The ROC curve generated for the PLS-DA model built in this study, based on internal validation, is seen in Figure 4. The most optimum threshold for discrimination in this study is designated by the point at (0,1), which corresponds to a cut-off value of 77%. This threshold indicates if 77% or more of the total spectra from a donor in the external validation dataset are assigned to the MDX class, than the overall prediction of the donor would be as belonging to the MDX class.

**Figure 4.**
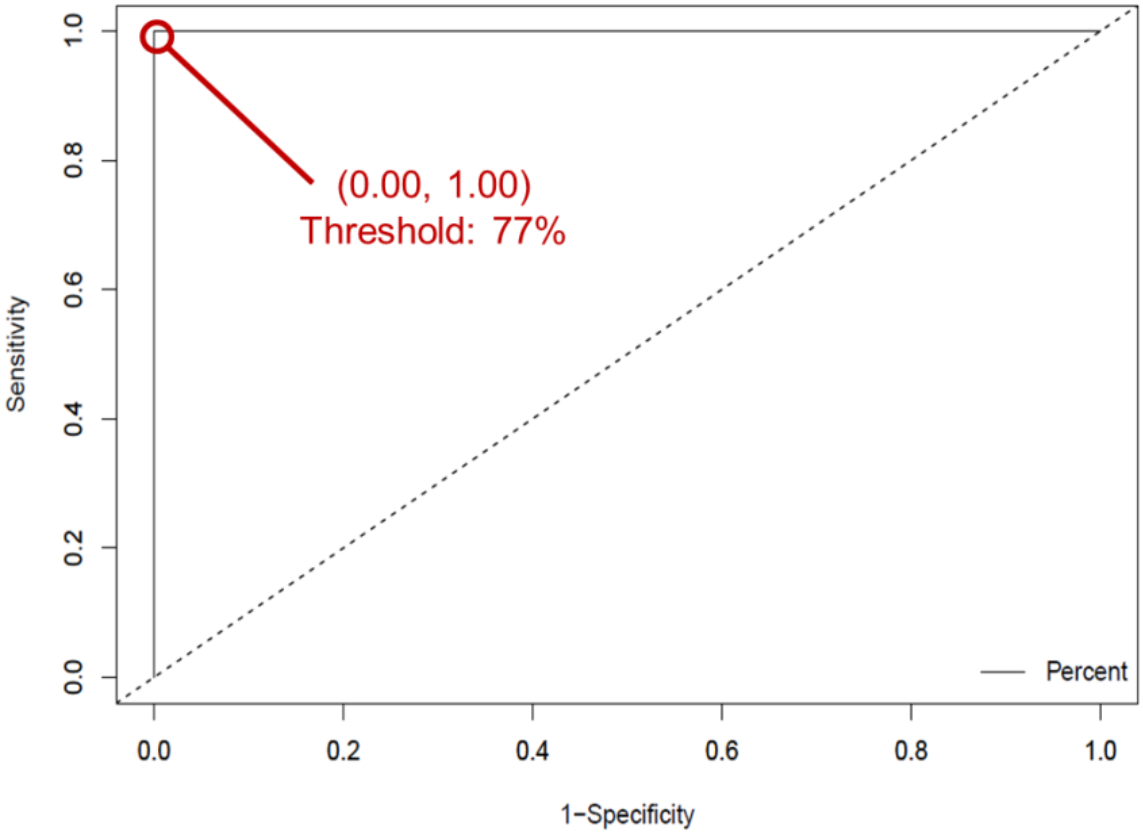
Receiver operating characteristic (ROC) curve. ROC curve for the internally cross-validated PLS-DA model, trained to differentiate between diseased and healthy control mice blood serum. The true positive rate (sensitivity) of each potential discrimination threshold are plotted according to each corresponding false positive rate (1 – specificity). The optimal threshold is designated by the point at (0.00, 1.00), corresponding to a threshold of 77%.

The threshold established by the ROC curve (77%) was applied to the model’s spectral-level predictions in order to generate a diagnosis at the donor level, as shown in Figure 5. The percentage of spectra which were identified as belonging to the MDX class is plotted. All four donors in the validation dataset were correctly identified. Thus, based on donor-level predictions, 100% successful external validation was achieved. This indicates the strength and capability of the model to be applied to new, unknown data, to make accurate diagnoses.

**Figure 5.**
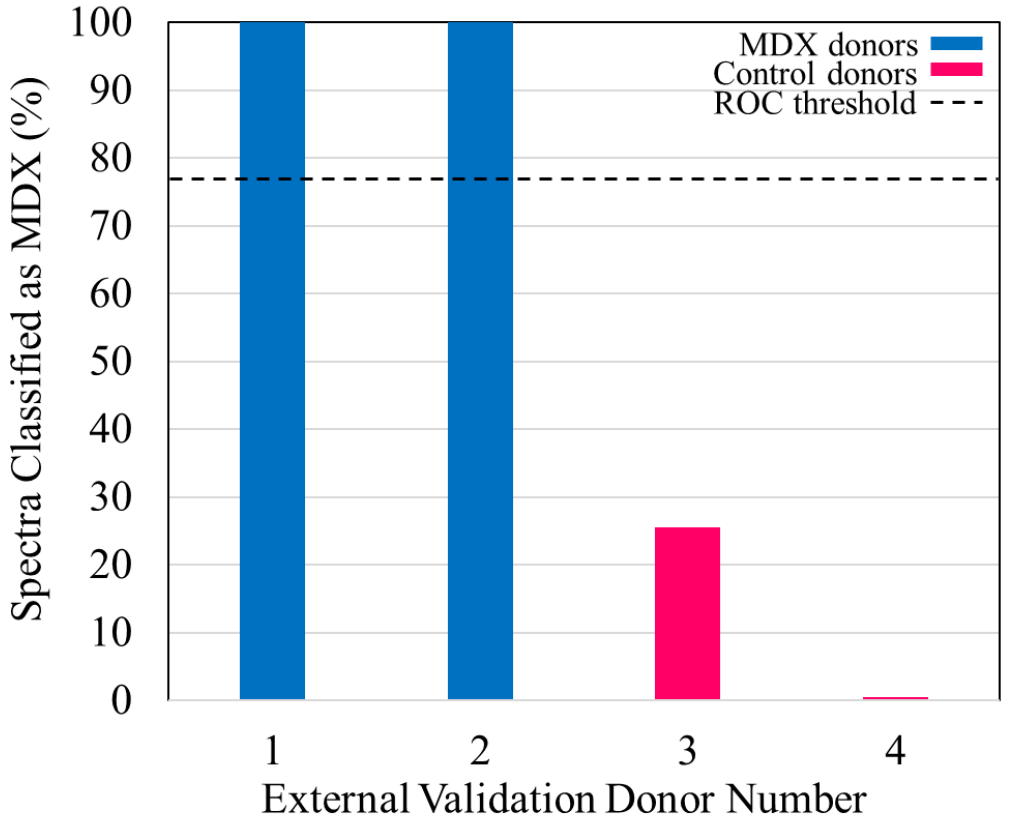
Histogram displaying the results of external validation of the PLS-DA model. The percentage of spectra classified as MDX is plotted as the bar height of each of the donors. The 77% threshold was established by the ROC curve and is plotted as the dashed line.

### Genetic Algorithm for identifying spectral differences in blood serum

In order to gain an understanding of the biochemical basis responsible for the model’s ability to discriminate between spectral datasets, Genetic Algorithm (GA) was performed. GA is a statistical technique which capitalizes on the ideas of “natural selection” and “survival of the fittest.”^25^ The algorithm will identify spectral features within the dataset which contribute the most discrimination power toward separating classes of data. The assignments of these selected spectral features provide an insight into the biochemical changes that occur as the disease progresses, and allow for the identification of key biochemical components which may serve as spectroscopic biomarkers for a disease. The results of GA are observed in Figure 6.

**Figure 6.**
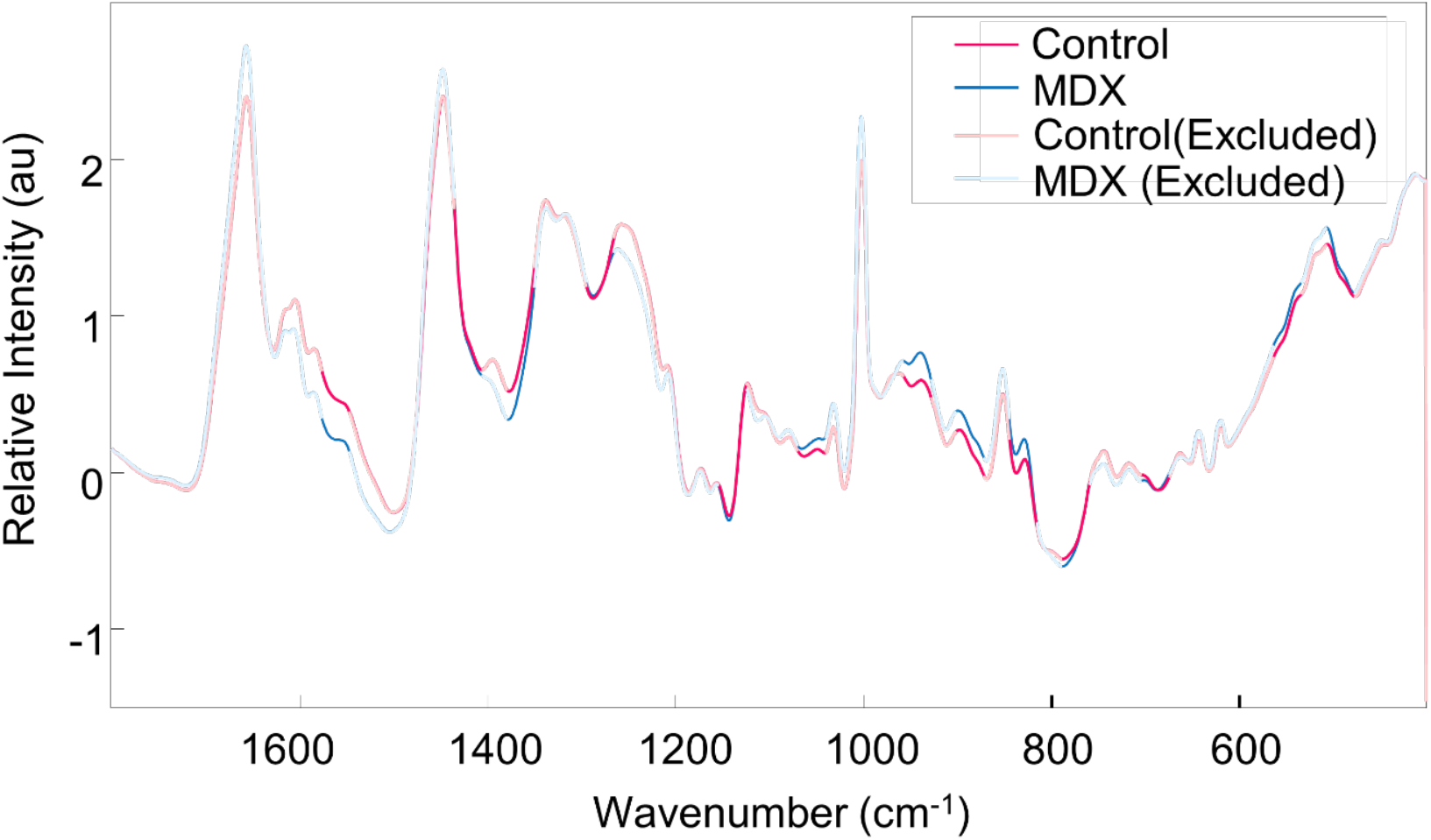
Genetic Algorithm analysis. Mean blood serum spectra of the two classes, including the spectral ranges selected by Genetic Algorithm: control (pink) and MDX (blue). Areas selected by Genetic Algorithm are marked by bolded lines. Spectral regions deemed as uninformative for discrimination are seen as unfilled lines.

The areas which are bolded in Figure 6 are those which were identified by GA as providing the most useful spectral information for discrimination purposes. Interestingly, the tentative assignment of these bands can be attributed to various biomarkers which have been previously shown to be linked to DMD and are summarized in Supplementary Information Table S.1. Monosaccharides and polysaccharides contribute to spectral bands at 899 and 940 cm^−1^.^26^ Specifically, bands at 1048 and 1124 cm^−1^ are attributed to vibrational modes of glycogen and glucose.^27,28^ Different vibrational modes of lipids are seen around the 878, 1124 and 1338 cm^−1^ selected regions.^27–29^ Proteins are represented by the spectral features around 1124, 1156, 1260, and 1554 cm^−1^ and collagen tentatively shows contributions at 507 and 1338 cm^−1^.^27–29^ Other GA-selected regions show potential influences from cholesterol (541, 702, and 959 cm^−1^), and DNA/RNA bases (750 and 829 cm^−1^).^26,27,29,30^ These spectral features, identified by GA, are assigned to major biological components which are associated with known biochemical changes that occur during the onset and progression of muscular dystrophy.

## Discussion

The combination of Raman hyperspectroscopy and advanced statistical analysis is incredibly advantageous for disease diagnostic purposes. Raman hyperspectroscopy involves the collection of multiple Raman spectra from a sample in order to characterize its heterogeneity and multicomponent composition. This is accomplished through acquiring the combination of spectral information alongside spatial information, allowing for the formation of a three dimensional data cube (*x, y, λ*). Two dimensions, *x* and *y*, correspond to spatial coordinates, and the third dimension, *λ*, represents the Raman spectrum collected at a particular pair of coordinates. By probing multiple small volumes or areas of a sample, there is a potential to identify biochemical components which, although may be present at low average concentrations, are present at a particular coordinate at a high local concentration. The ability to detect such components using this method indicates that they may be useful for discrimination purposes, and can serve as spectroscopic biomarkers. Thus, the advantage of Raman hyperspectroscopy resides in its ability to detect multiple biomarkers simultaneously, which can potentially be used for discrimination and diagnostic purposes.^16^

It is often observed that spectral differences between two similar classes of samples, such as healthy and diseased body fluids, are insignificant when evaluated at the average level.^17,31^ It is expected that the majority of the composition of a body fluid remains consistent between healthy and diseased donors. In this research, the difference spectrum calculated between the average control spectrum for all control donors and the average MDX spectrum of all MDX donors was shown to be statistically insignificant. This is an indication that statistical analysis is required to better understand and evaluate the Raman spectral data obtained, and specifically, to uncover hidden characteristic features of the two classes as well as spectral variability which can be capitalized on for building a discrimination algorithm. In this study, the combination of Raman hyperspectroscopy and advanced statistical analysis was used to develop an algorithm which could accurately identify Duchenne muscular dystrophy via mice blood serum.

The *mdx* mice model was specifically selected for this project because the species exhibits a mutation within its DMD gene which results in the mouse not producing a functional dystrophin protein and thus developing the disease. This animal model has been widely studied in the last several decades, and has provided extensive insight into the pathophysiology associated with muscular dystrophy.^2,13^ Additionally, the *mdx* mouse model can be manipulated to test potential therapeutic strategies, and lack of interfering factors, such as comorbidities or influence of prescribed medications, makes it ideal for evaluating novel diagnostic methods. As such, this model was selected for the initial proof-of-concept diagnostic study, which can further be easily translated into a platform for larger and more advanced diagnostic studies within humans.

Two different statistical techniques were used in this study. The first, PLS-DA, was selected to generate the prediction algorithm. The 14 donors used in this study were split into two groups: a calibration set and a validation set. The spectral data from the calibration set, consisting of 452 total spectra from five control donors and five MDX donors, was used to build and train the prediction algorithm. Internal validation by venetian blinds resulted in 95.2% sensitivity and 94.6% specificity for identifying MDX spectra as compared to control spectra within the calibration dataset.

The prediction capabilities of the algorithm was then tested through external validation using the validation set of samples, containing two control donors and two MDX donors. The spectral data from these four samples was used to test the ability of the algorithm to make predictions regarding samples it has never before seen, and thus cannot have an inherent bias toward. The PLS-DA algorithm generated classification predictions for each individual spectrum collected from the four donors. Each sample is represented by a multitude of spectra; because blood serum is inherently heterogeneous, each spectrum is expected to deviate from the mean to some extent. It is also expected that a portion of the mice blood serum components are the same between control and *mdx* donors. As such, it is reasonable to assume that some spectra from one class may be predicted as belonging to the other, due to the natural overlap in biochemical composition. In order to better translate individual spectral predictions to an over-all prediction for the donor, ROC curve analysis was used to establish an optimum threshold, or cut-off, for donor-level predictions. Using the determined threshold of 77%, all four donors of the validation dataset were identified as belonging to their true class. External validation is an established process for determining whether or not a model is robust enough for successful application to new, and unknown, spectral data for accurate predictions.^32,33^ Thus, successful external validation, as achieved here, indicates the potential for the method to be applied within diagnostic settings.

Following external validation, an attempt was made to understand the major biochemical differences which are useful for distinguishing between the classes of Raman spectral data. Past literature has demonstrated strong links between the pathogenesis of DMD and the aforementioned biomolecules (Supplementary Information Table S.1). Specifically, studies have shown that a general increase in lipids, including triglycerides, phospholipids, cholesterol, and cholesterol esters, is found in patients with muscular dystrophy.^34,35^ In fact, in *mdx* mice, elevated lipid levels were found to be associated with significant exacerbation of muscle pathology, including myofiber damage and skeletal muscle remodeling.^35^ Collagen has also been found to play a role in the pathogenesis of muscular dystrophy.^36^ Among the evidence, researchers found an inverse relationship exists between the over-production of connective tissue and muscle protein synthesis in patients suffering from DMD.^37–39^ Other research observed unusual clusters of “sticky cells” formed by dissociated muscle of patients with Duchenne and Becker muscular dystrophies, a sign which reflects abnormal collagen production.^40^ Mutations in genes coding for collagen type VI are also responsible for congenital muscular dystrophies including Bethlem myopathy and Ullrich congenital muscular dystrophy.^41^

Many serum proteins have been identified as biomarkers which reflect the pathogenesis of DMD; the concentration of 23 identified mouse serum proteins exhibited an increase while four other proteins were found to exist at concentrations significantly lower in *mdx* mice as compared to healthy control mice in one study. Proteins which were elevated mostly originated from muscle or were glycolytic enzymes, transport proteins, or other proteins such as creatine kinase M.^42^ These identified protein biomarkers reflect the muscle activity as well as pathogenesis of the disease.

Many more studies have also identified various serum proteins as biomarkers for muscular dystrophy.^43–46^ It is thus unsurprising that GA identified spectral features which can be attributed to vibrational modes of proteins as being useful for discrimination purposes. Furthermore, a relationship between glycogen metabolism and DMD was supported by Naim et al. Here, results show that *mdx* mice have increased skeletal muscle glycogen content; many of the enzymes involved in the skeletal muscle glycogen metabolism were dysregulated.^47^ Because of the dysregulation of glycogen, levels of glucose in the blood may be affected, connecting the identification of both glycogen and glucose here as also being important spectroscopic markers for DMD.

Notably, the spectral features identified by GA as being the most useful for spectroscopically discriminating between the two classes of data can also be assigned to vibrational modes of classes of biomolecules which have previously been related to the pathogenesis of the disease itself. Clearly, there is a connection between the progression of the disease and the spectroscopic signature produced. This link is strong enough to provide identifiable information which can be capitalized on through advanced statistical analysis for the purpose of generating a successful diagnostic algorithm.

The contribution of multiple biomarkers to the spectroscopic signature of DMD allows for much more specific identification of the disease, and further supports the strength of the method. In general, by identifying biochemical components whose alterations in composition or concentration reflect the stage of a particular disease, the ability to detect that disease is dramatically increased, and thus can result in very high levels of classification accuracy.^16^ Through the identification of the aforementioned biomolecules associated with DMD, we were indeed able to achieve high levels of diagnostic accuracy. Raman hyperspectroscopy allows for simultaneous detection of multiple, potentially new, biomarkers for a disease, as is observed herein. This is incredibly advantageous over other diagnostic methods which simply investigate one, known, biomarker at a time.

The method of combining Raman hyperspectroscopy with advanced statistical analysis is shown in this proof-of-concept study to be successful for detecting Duchenne muscular dystrophy. Raman spectra were collected from the blood serum of mice who were either healthy controls or who had the disease. The spectral data was analyzed using PLS-DA, which showed 95.2% sensitivity and 94.6% specificity for identifying MDX spectra in the calibration dataset, and 100% sensitivity and 87.0% specificity for identifying MDX spectra in the validation dataset. Based on donor-level predictions generated using ROC curve analysis, 100% accuracy was achieved for correctly predicting to which class the donors in the external validation dataset belonged. This is the first time this methodology has been applied toward diagnosing DMD. Further, Genetic Algorithm identified key biochemical components which were responsible for spectroscopic discrimination, indicating a link between the disease progression and the Raman spectroscopic fingerprint produced. Future research is required to study this link on a larger scale, and to investigate if a similar trend is observed within humans. However, it is clear that this methodology has significant potential for use as a novel technique for diagnosing Duchenne muscular dystrophy.

## Methods

### Mouse strains and sample collection

The *mdx* (C57BL/10ScSn-Dmd<mdx>/J; Stock Number 001801) and counterpart control mice (C57BL/10ScSnJ; Stock Number 000476) were purchased from the Jackson Laboratory, Bar Harbor, ME, USA. The mice were raised following the protocol approved by the Institutional Animal Care and Use Committee to the appropriate age (3 months and 12 months) before harvesting the tissue and blood samples. As *mdx* is an X-linked muscle degenerative disease, male *mdx* and male control mice were studied.

Mice were euthanized following the standard operating procedure of Laboratory Animal Resources (LAR SOP # 105 and 106). Briefly, the mice were first anesthetized to a surgical plane of anesthesia under isoflurane inhalation using an induction chamber. The depth of anesthesia was verified by establishing the loss of pedal reflex. The mice were then euthanized under anesthesia by isoflurane and then by cervical dislocation. For harvesting skeletal muscle, the hind leg skins were removed and the Tibialis Anterior (TA) muscles were removed by a surgical blade. The TA muscles were cut into 2 pieces and frozen fresh with Optimal Cutting Temperature (OCT) compound in plastic molds. The freezing process was carried out in a jar containing semi-frozen iso-butanol and again frozen in liquid nitrogen before storing the tissue blocks at −80°C. The blood samples were collected from the euthanized mice by cardiac puncture. Briefly, the skin and the rib cases were cut and pinned in the dissection board. The jugular vein was cut by sharp scissors and blood was collected in small Eppendorf tubes, without use of anticoagulant, using pasteur pipettes.

The used gloves, bench coats, and paper towels were collected in the biohazard containers and surgical blades were collected in the biohazard labeled sharp container and discarded following the instructions of the Institutional Biosafety Committee. The experimental bench was cleaned first with 50% beach, then with sterile water, and finally with 70% ethanol.

### Isolation of serum

The serum was isolated following a standard laboratory protocol. Briefly, the tubes containing the blood without any anticoagulant were left at room temperature in a standing position for about 35 minutes, allowing the blood to clot. Then, the clotted blood samples were centrifuged at 20°C and 2000g for 15 minutes; the serum fraction was moved to a fresh tube and stored at −80°C. At the time of analysis, the blood serum was allowed to thaw. Each serum sample (10 μL) was deposited on an aluminum foil substrate and set aside to dry overnight before analysis.

### Cryosection and histochemistry of TA muscle

The cryosections and H&E staining was carried out using established protocol as described elsewhere.^48,49^

### Raman hyperspectroscopic methods

A Renishaw inVia Raman spectrometer equipped with a research-grade Leica microscope was used to collect Raman spectra. A PRIOR automatic mapping stage was used during measurements and the 50X objective was used to focus on the sample. Spectra were recorded between 400-1800 cm^−1^ under excitation by the 785 nm diode laser, which was reduced to about 50% laser power to prevent photo-degradation of the sample. For each sample, 50 spectra were recorded to capture the inherent heterogeneity of blood serum.

### Data treatment and advanced statistical analysis

Spectra were recorded using WiRE 3.2 software, and then imported to MATLAB version 2017b workspace (Mathworks, Inc.). Any individual Raman spectrum which displayed a poor signal-to-noise ratio or exhibited cosmic rays was removed from the dataset. The remaining spectra were subjected to preprocessing.

#### Partial Least Squares Discriminant Analysis (PLS-DA)

PLS Toolbox (eigenvector Research, Inc.) was used for statistical analysis. PLS-DA was selected to accomplish discrimination between the healthy and diseased classes. PLS-DA algorithms have been shown to be effective in various disease diagnostic applications including for investigating inflammatory bowel diseases^50^, coronary heart diseases^51^, and various forms of cancer^52–64^, among many others. Specifically, PLS-DA is a supervised technique which is used to predict categorical variables. The dataset being analyzed is reduced to a few latent variables (LVs), which capture the maximum covariance between spectral data and the labeled classes. Each spectrum is then given a score which corresponds to how closely that spectrum resembles a particular LV. Different classes of samples will be represented by a set of scores seen as characteristic for a sample within that class.^65^ In this way, unknown samples can be identified through comparison of the unknown sample’s score to those of classes which are known. Here, PLS-DA was built using spectral data from ten samples (five control, five MDX); eight LVs were used to reduce the dimensionality of the dataset. The performance of the algorithm was investigated using venetian blind internal validation. Following internal validation, predictions of unknowns were made using the spectral data obtained from donors of the external validation dataset.

#### Genetic Algorithm (GA)

GA was used to determine the spectral features which were the most useful for discrimination between the two classes of data. GA is a statistical technique inspired by the ideas of evolution. The algorithm aims to solve a specific problem by generating potential solutions; recombination operators are applied to the data in order to preserve critical information which can best solve the problem.^66^ Essentially, GA will identify spectral variables which provide the lowest prediction error rates, identified through a repetitive algorithm building process. In this way, it can recognize which spectral features of the dataset provide the most useful information for discriminating between different classes of data. Concurrently, it will eliminate uninformative data as well as noise from future consideration. Here, GA was applied to the training dataset which consisted of ten donors and 452 spectra. The parameters of GA are given as follows: the population size was set to 80; the mutation rate to 0.005, and the maximum number of generations for each run to 100. The breeding was fixed to double crossover, the window width was 30, and 30% of the windows were initially included. To identify the diagnostic features from within the measured Raman spectral dataset, GA was independently run 100 times which allowed for identification of significant spectral bands useful for discrimination purposes. The identified spectral features were assigned to tentative corresponding vibrational modes, according to the literature, to determine potential biochemical basis responsible for spectroscopic differentiation (Supplementary Information Table S.1).

## Data Availability

The data that support the findings of this study are available from the corresponding author upon reasonable request.

## Acknowledgements

This work was supported by the SUNY startup and American Heart Association (AHA 17SDG33670339) grants to B.K.D. N.M.R was supported by NIH training grant T32 GM13206.

## Author contributions

I.K.L and B.K.D conceived the project; N.M.R, B.K.D. and I.K.L contributed to study design; N.M.R. and A.G. contributed to data collection; P.D. was involved in mouse colony maintenance, harvesting blood and tissues, histology, validation of mdx model system, and isolation of blood serum and writing the relevant part of the manuscript; I.K.L and B.K.D. provided expertise with data interpretation and supervision of the project.

## Supplementary Information

**Figure S.1.**
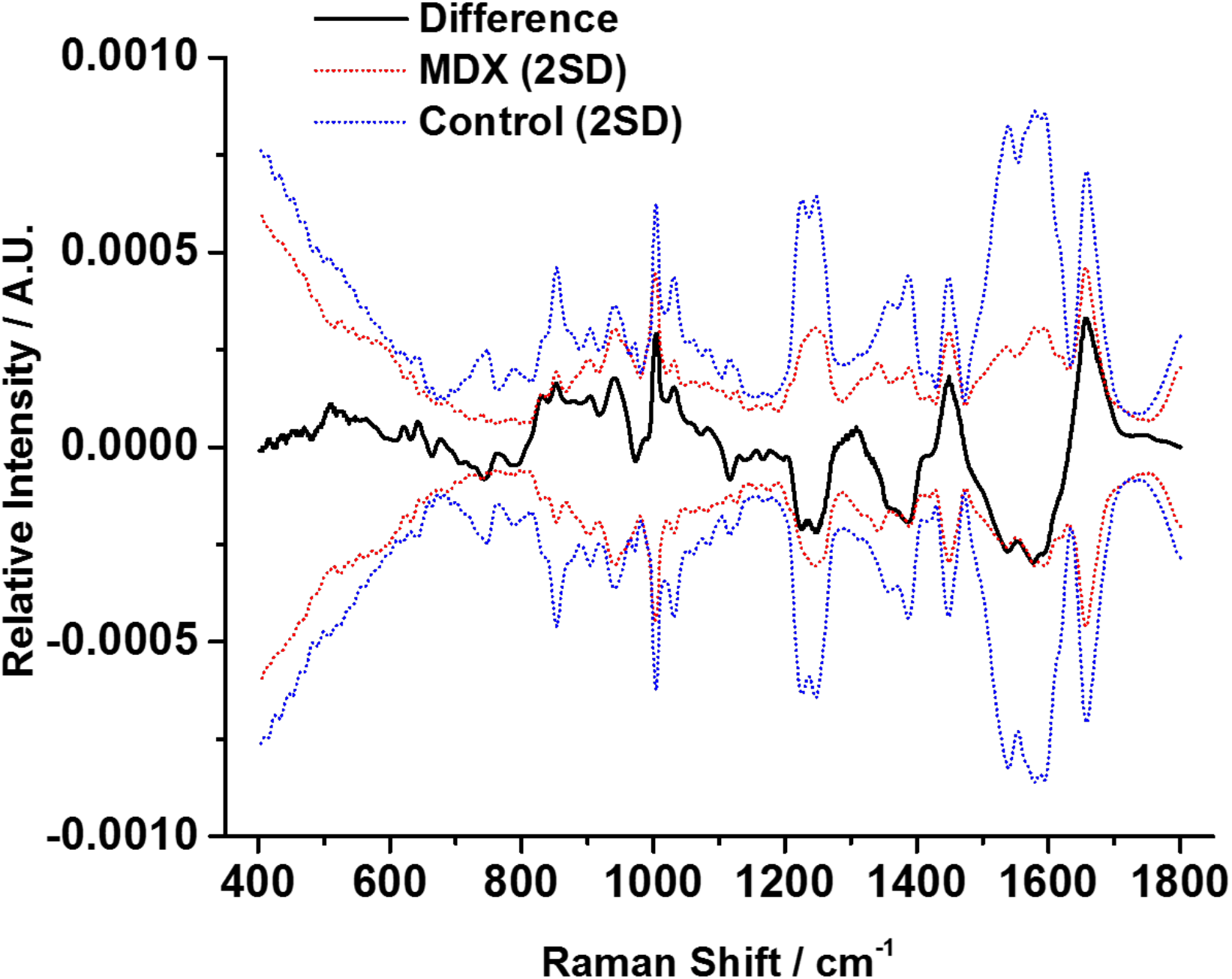
The difference between the average Control and average MDX spectra. The pre-processed difference mean blood serum spectra between Control and MDX (bold line) with ± 2 standard deviations (thin dotted lines) of the Control (blue) and MDX (red) donors’ spectral data sets.

**Table S.1.** Tentative assignments of the most important regions in the Raman spectrum of blood serum for discrimination between HC and MDX mice, as determined by GA.

|  | **<u>GA</u>**<br>**<u>region</u>** | **<u>Peak Position</u>**<br>**<u>(cm<sup>-1</sup>)</u>** | **<u>Vibrational Mode</u>** | **<u>Contributions<sup>26-30</sup></u>** |
| --- | --- | --- | --- | --- |
| 1 | 479-507 | 507 | (S-S)<sup>a</sup> | Collagen; Cysteine |
| 2 | 535-563 | 541 | (S-S)<sup>a</sup> | Cysteine; Cholesterol |
| 3 | 675-703 | 702 |  | Cholesterol; Cholesterol Ester |
| 4 | 760-786 | 750 | Ring breathing mode | Pyrimidines of DNA/RNA bases |
| 5 | 815-844 | 829 | (O-P-O)<sup>a</sup>; out-of-plane<br>ring breathing | DNA/RNA; Tyrosine |
| 6 | 872-899 | 878 | (C-C-N<sup>+</sup>)<sup>b</sup> | Lipids |
|  |  | 899 | (C-O-C) skeletal mode | Monosaccharides; Disaccharides |
| 7 | 929-960 | 940 | Skeletal mode | Polysaccharides |
|  |  | 959 |  | Cholesterol |
| 8 | 1042-1066 | 1048 |  | Glycogen |
| 9 | 1124-1156 | 1124 | (C-C)<sup>a</sup>; (C-N)<sup>a</sup> | Lipids; Proteins; Glucose |
|  |  | 1156 | (C-C)<sup>a</sup> | Proteins |
| 10 | 1260-1294 | 1260 | Amide III | Proteins |
| 11 | 1340-1377 | 1338 | (CH<sub>2</sub>/CH<sub>3</sub>)<sup>c,d</sup> | Collagen; Lipids |
| 12 | 1547-1575 | 1554 | Amide II | Proteins |

<sup>a</sup>stretching; <sup>b</sup>symmetric stretching; <sup>c</sup>wagging; <sup>d</sup>twisting

